# Schistosomes alter expression of immunomodulatory gene products following *in vivo* praziquantel exposure

**DOI:** 10.1101/2020.01.22.915710

**Authors:** Paul McCusker, Claudia M. Rohr, John D. Chan

## Abstract

Control of the neglected tropical disease schistosomiasis relies almost entirely on praziquantel (PZQ) monotherapy. How PZQ clears parasite infections remains poorly understood. Many studies have examined the effects of PZQ on worms cultured *in vitro*, observing outcomes such as muscle contraction. However, conditions worms are exposed to *in vivo* may vary considerably from *in vitro* experiments given the short half-life of PZQ and the importance of host immune system engagement for drug efficacy in animal models. Here, we investigated the effects of *in vivo* PZQ exposure on *Schistosoma mansoni*. Measurement of pro-apoptotic caspase activation revealed that worm death occurs only after parasites shift from the mesenteric vasculature to the liver, peaking 24 hours after drug treatment. This indicates that PZQ is not directly schistocidal, since the drug’s half-life is ∼2 hours, and focuses attention on parasite interactions with the host immune system following the shift of worms to the liver. RNA-Seq of worms harvested from mouse livers following sub-lethal PZQ treatment revealed drug-evoked changes in the expression of putative immunomodulatory and anticoagulant gene products. Several of these gene products localized to the schistosome esophagus and may be secreted into the host circulation. These include several Kunitz-type protease inhibitors, which are also found in the secretomes of other blood feeding animals. These transcriptional changes may reflect mechanisms of parasite immune-evasion in response to chemotherapy, given the role of complement-mediated attack and the host innate / humoral immune response in parasite elimination. One of these isoforms, SmKI-1, has been shown to exhibit immunomodulatory and anti-coagulant properties. These data provide insight into the effect of *in vivo* PZQ exposure on *S. mansoni*, and the transcriptional response of parasites to the stress of chemotherapy.

**Author Summary:** The disease schistosomiasis is caused by parasitic worms that live within the circulatory system. While this disease infects over 200 million people worldwide, treatment relies almost entirely on one drug, praziquantel, whose mechanism is poorly understood. In this study, we analyzed the effects of praziquantel treatment on the gene expression of parasites harvested from mice treated with praziquantel chemotherapy. Despite the rapid action of the drug on worms *in vitro*, we found that key outcomes *in vivo* (measurement of cell death and changes in gene expression) occurred relatively late (12+ hours after drug administration). We found that worms increased the expression of immunomodulatory gene products in response to praziquantel, including a Kunitz-type protease inhibitor that localized to the worm esophagus and may be secreted to the external host environment. These are an intriguing class of proteins, because they display anti-coagulant and immunomodulatory properties. Up-regulation of these gene products may reflect a parasite mechanism of immune-evasion in response to chemotherapy. This research provides insight into the mechanism of praziquantel by observing the effect of this drug on worms within the context of the host immune system.

## Introduction

The neglected tropical disease schistosomiasis is caused by infection with parasitic *Schistosoma* blood-flukes and afflicts over 200 million people worldwide. These parasites can survive for years – even decades – within the host circulatory system, employing various mechanisms including mimicry of host glycans [1], binding non-immune immunoglobulins [2], and secretion of immunomodulatory extracellular vesicles [3]. With no vaccine available, control of this disease is almost entirely reliant upon chemotherapy with one drug – praziquantel (PZQ)[4].

PZQ’s anti-parasitic mechanism of action remains poorly understood. Since the initial studies on this drug over four decades ago, it has been clear that a hallmark of PZQ action on worms is rapid, Ca^2+^-dependent contractile paralysis [5], with several Ca^2+^ channels having been proposed as the drug’s target(s) [6, 7]. However, while parasite contraction provides a clear visual readout of drug action *in vitro*, the mechanism of PZQ-evoked parasite elimination *in vivo* is more complex. For example, PZQ causes contractile paralysis of both immature and sexually mature worms in vitro, despite the fact that only sexually mature worms and not the immature liver-stage parasites are susceptible to PZQ treatment *in vivo* (Xiao et al., 1985). Whether PZQ-evoked contraction is related to tegument damage, the other signature effect of anthelmintic exposure, is also unclear. Muscle contraction occurs within seconds, but tegument depolarization occurs over a period of several minutes [8], and pharmacological experiments do not show a correlation between these two phenotypes [9]. Therefore, the *in vitro* phenotype of worm contraction, while useful for drug screening, provides an incomplete readout of PZQ’s mechanism of action, which likely encompasses a cascade of events that trigger immune-mediated elimination of the parasites *in vivo*.

PZQ efficacy *in vivo* requires engagement of the host immune system. Following PZQ exposure, sexually mature worms display damage to the tegument surface, which exposes parasite antigens to the host humoral immune system [10] and triggers the recruitment of innate immune cells likely involved in parasite elimination [11]. PZQ is less efficacious at clearing infections from immunocompromised models such as T-cell [12] and B-cell deprived mice [13]. A requirement for the host immune system may also contribute to PZQ’s lack of efficacy against immature parasites, since only after worms become sexually mature and begin egg laying does the host respond with a wave of macrophage recruitment to the liver and an acute Th2 response [14, 15]. Notably, PZQ is ineffective against unisex female infections, which do not reach sexual maturity and do not lay eggs [16].

Given the importance of the host immune system to PZQ action, we sought to characterize *Schistosoma mansoni* transcriptional changes following *in vivo* drug exposure. Prior microarray experiments based on expressed sequence tag (EST) libraries have investigated changes in gene expression following *in vitro* PZQ treatment of *S. mansoni* [17] or *in vivo* PZQ treatment of *Schistosoma japonicum* [18]. But no comprehensive study of genome-wide changes in gene expression following in vivo PZQ treatment has been performed in the decade since these parasites’ genomes have been sequenced. We established conditions for PZQ dosing that elicited a sublethal response in parasites *in vivo*, sequenced the transcriptomes of both sexually mature (7-week-old) and immature (4-week-old) infections treated with either vehicle control or PZQ, and then mapped the expression patterns of differentially expressed transcripts by *in situ* hybridization. These data revealed that numerous up-regulated transcripts were expressed near the esophagus – a location previously identified as a hotspot for expression of immunomodulatory gene products. Specifically, these data highlight a clade of up-regulated Kunitz-type protease inhibitors. Given the immunomodulatory and anti-coagulant activity reported for one of these isoforms, SmKI-1 [19], these changes may reflect the response of parasites to the hostile, immune cell-rich environment of the liver following PZQ treatment.

## Materials and Methods

Ethics statement. Animal work was carried out with the oversight and approval of the Laboratory Animal Resources facility at the Medical College of Wisconsin, adhering to the humane standards for the health and welfare of animals used for biomedical purposes defined by the Animal Welfare Act and the Health Research Extension Act. Experiments were approved by the Medical College of Wisconsin IACUC committee (approved protocol numbers AUA00006471 and AUA00006735).

*In vitro* schistosome assays. Female Swiss Webster mice infected with *S. mansoni* cercariae (NMRI strain) were sacrificed by CO_2_ euthanasia at 4 weeks (for immature worms) or at 7 weeks post-infection (for mature worms). Immature worms were recovered from mouse livers, and mature worms were recovered from the mesenteric vasculature. Harvested worms were washed in DMEM (ThermoFisher cat. # 11995123) supplemented with HEPES (25mM), 5% v/v heat inactivated FCS (Sigma Aldrich cat. # 12133C) and Penicillin-Streptomycin (100 units/mL). Worms were cultured in 6 well dishes (4-5 mature male worms or 8-10 immature worms in 3mL media per well) at varying concentrations of praziquantel (PZQ, Sigma-Aldrich cat. # P4668) or DMSO vehicle control and imaged to record phenotypes.

Cell proliferation assay. Immature and mature worms (harvested 25 and 49 days post-infection) were treated with drug (37°C, 14 hours). Worms were washed in drug-free media and allowed to recover for 8 hours, before media was then supplemented with EdU (ThermoFisher Scientific cat. # C10637, 10μM) for a further 14 hours. Worms were fixed in 4% PFA in PBST (PBS + 0.3% triton X-100), washed in PBST, followed by 1:1 PBST:MeOH, and stored in 100% MeOH at −20°C. Worms were rehydrated in 1:1 PBST:MeOH, bleached (5% formamide, 0.5X SSC and 1.2% hydrogen peroxide in ddH_2_O), rinsed in PBST and permeabilized (0.1% SDS and 0.01mg/ml proteinase K in PBST) for either 30 min (7-week-old schistosomes) or 15 min (4-week-old schistosomes), post-fixed (4% PFA, 10 min), and EdU detection was performed using 1mM CuSO_4_, 0.1mM Azide-fluor 488 (ThermoFisher Scientific cat. # 760765) and 100mM ascorbic acid in PBS. Worms were stained with DAPI (1µg/ml) and loaded into a 96 well optical bottom black plate for imaging using the ImageXpress Micro Confocal system (Molecular Devices).

*In vivo* hepatic shift assay. Mice harboring mature infections (6-7 weeks) were administered a fully curative single dose of PZQ (400 mg/kg PZQ dissolved in vegetable oil and delivered by oral gavage) or a sub-curative dose of PZQ (100 mg/kg PZQ solubilized in 50 µL DMSO, then diluted in 200 µL 5% w/v Trappsol (Cyclodextrin Technologies Development cat. # THPB-p-31g) in saline (NaCl 0.9%) solution and delivered by intraperitoneal injection). Mice were sacrificed by CO_2_ euthanasia at varying timepoints after drug administration and worms were recovered from either the mesenteries, portal vein or liver. Data = mean ± standard error for 3-5 mice per cohort.

Measurement of caspase 3/7 activation. Pro-apoptotic caspase 3/7 activation was measured in worms harvested from mice following drug treatment using the Caspase-Glo 3/7 Assay Kit (Promega). Worms were harvested from either the mesenteries or liver of mice, then homogenized in assay buffer (PBST, supplemented with HEPES 10mM and protease inhibitor (Roche cOmplete Mini EDTA-free Protease Inhibitor Cocktail)) using a mini mortar and pestle and stored at −80°C. Worm homogenate (5 pooled male and female worm pairs / 125μL assay buffer) was diluted 1:5 in distilled water and added to Caspase-Glo 3/7 substrate (1:1 volume ratio) in solid white 96 well plates. Luminescence was read using a SpectraMax i3x Multi-Mode Microplate Reader (Molecular Devices). Data reflect mean ± standard error of ≥ 3 biological replicates.

Transmission electron microscopy. Worms were harvested from infected mice and treated with PZQ as described for movement assays, then fixed overnight at 4°C in 2.5% glutaraldehyde / 2% paraformaldehyde in 0.1M sodium cacodylate (pH 7.3). Worms were then washed in 0.1M sodium cacodylate (3×10 minutes) and post-fixed on ice (2 hours) in reduced 1% osmium tetroxide. Worms were washed in distilled water (2×10 minutes), stained in alcoholic uranyl acetate (overnight, 4°C), rinsed in distilled water, dehydrated in MeOH (50%, 75% and 95%), followed by successive rinses (10 minutes) in 100% MeOH and acetonitrile. Worms were incubated in a 1:1 mixture of acetonitrile and epoxy resin for 1 hour prior to 2×1-hour incubations in epoxy resin, then cut transversely and embedded overnight in epoxy resin (60°C). Ultra-thin sections (70nm) were cut onto bare 200-mesh copper grids, stained in aqueous lead citrate (1 minute), then imaged on a Hitachi H-600 electron microscope fitted with a Hamamatsu C4742-95 digital camera operating at an accelerating voltage of 75 kV.

Comparative RNA-Seq. For experiments in Figure 1, mice were treated with a single, curative dose of PZQ (400 mg/kg) delivered by oral gavage, sacrificed by CO_2_ euthanasia at various time-points, and worms were harvested from either the mesenteries or liver and homogenized in Trizol Reagent (Invitrogen). For experiments on mice treated with a sub-lethal dose of PZQ (100 mg/kg by intraperitoneal injection) at 4-weeks or 7-weeks post-infection, animals were sacrificed 14 hours later and worms were harvested. For 4-week-old infections, worms were harvested from the livers of vehicle control and PZQ treated mice. For 7-week-old infections, worms were harvested from either the mesenteric vasculature (vehicle control cohort) or the liver (PZQ treated cohort). Parasites were homogenized in Trizol, and libraries were generated using the TruSeq Stranded mRNA kit (Illumina), sequenced using the Illumina HiSeq 2500 system (high-output mode, 50 bp paired-end reads at 20 million reads per sample), and trimmed reads were mapped to the *Schistosoma mansoni* genome (v7.2) using HISAT2. Differentially expressed gene products between vehicle control and PZQ-treated samples were identified using EdgeR (tagwise dispersion model, FDR adjusted p-value < 0.05). Differentially expressed, up and down-regulated transcripts were ranked by fold change and then functional enrichment analysis was performed using g:Profiler [20] to identify enriched GO-terms and KEGG pathways. Read counts and differential expression data are contained in S1-S3 Files. FASTQ files containing RNA-Seq data have been deposited in the NCBI SRA database under accession numbers PRJNA597909 and PRJNA602528.

**Figure 1.**
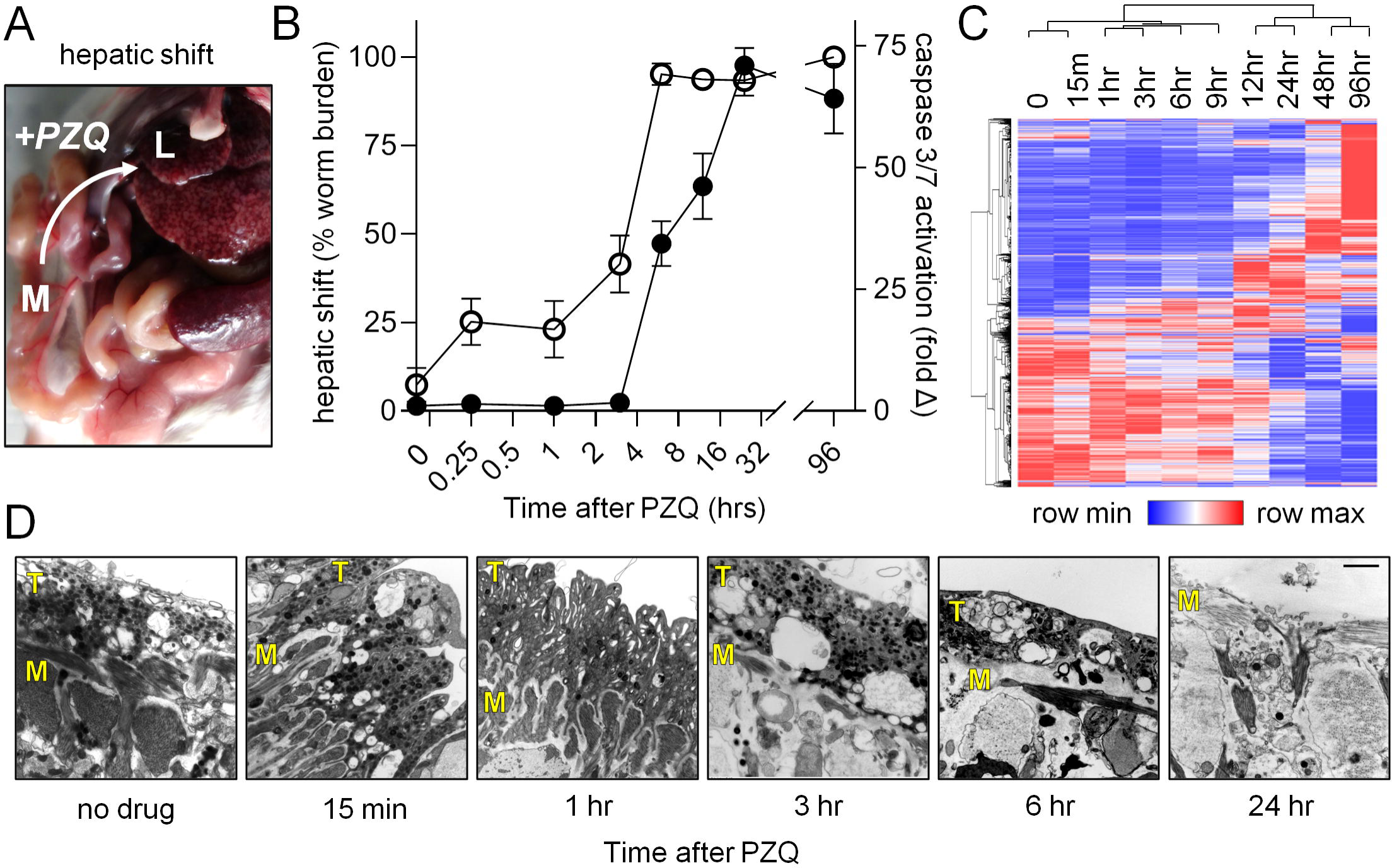
Parasite death occurs following *in vivo* hepatic shift. **(A)** A curative dose of PZQ (400 mg/kg) was administered to mice 7 weeks post-infection and worms were harvested at various time points from either the mesenteries (M) or the liver (L). **(B)** Time course of parasite hepatic shift (open symbols, left axis) and pro-apoptotic caspase-3/7 activation (solid symbols, right axis). **(C)** Changes in gene expression in worms harvested at various timepoints following PZQ treatment in B. Heatmap reflects minimum (blue) and maximum (red) z-score values for all transcripts showing >1 log_2_ fold change and average TPM > 3 (see S1 File for raw data). **(D)** Transmission electron microscope (TEM) images of the dorsal male body wall showing the time course of PZQ-evoked tissue damage. T = tegument. M = muscle.

Molecular cloning. RNA was recovered from control worms, or those treated with sublethal dose of PZQ (100mg/kg), using the Purelink RNA Mini kit (ThermoFisher Scientific), with on-column DNase treatment. cDNA was synthesized using the High-Capacity RNA to cDNA kit (ThermoFisher Scientific). Transcripts were amplified using FastStart Taq DNA Polymerase Kit (Millipore Sigma) using primers (S4 File). Amplicons were ligated into pGEM T-easy vector (Promega) and Sanger sequenced.

*In situ* hybridization. Forward (control) and reverse (target) RNA probes were synthesized from plasmids amplified via PCR (Advantage HD Polymerase Kit, Takara Bio) using T7 or SP6 RNA Polymerases (ThermoFisher Scientific) along with DIG-UTP-labelling mix (Millipore Sigma). Worms harvested from the mesenteries of untreated mice were used to visualize transcripts Smp_076320, Smp_195070, Smp_200150, Smp_246770, Smp_302320, and Smp_311670. Transcripts Smp_008660, Smp_214060 and Smp_336990 were visualized in worms harvested from livers of mice treated with sublethal dose of PZQ (100mg/kg), due to low levels of expression in untreated worms. *In situ* was performed as per reference [21]. Recovered worms were relaxed in 0.25% tricane (2-3 mins), killed in 0.6M MgCl2, and fixed (4% PFA, overnight at 4°C). Worms were washed in PBST, 1:1 PBST:MeOH, and stored in 100% MeOH, −20°C. Worms were rehydrated in 1:1 PBST:MeOH, PBST, washed in 1X SSC (10 min), bleached (5% formamide, 0.5X SSC and 1.2% hydrogen peroxide in ddH2O), permeabilized (0.1% SDS and 0.01mg/ml proteinase K in PBST), post-fixed (4% PFA), wasjed in 1:1 PBST + prehybridization solution, and then incubated in prehybridization solution (2 hours, 52°C). Worms were treated with probes for ≥16 hours at 52°C, washed in dilutions of SSC (2x and 0.2x) and TNT prior to blocking (1-2 hours in blocking solution of 5% heat-inactivated horse serum (Millipore Sigma), 0.5% Western Blocking Regent (Millipore Sigma), and incubated overnight in anti-DIG-AP (1:2000, Millipore Sigma). Worms were washed (TNTx − 0.1 M Tris pH 7.5, 0.15 M NaCl, and 0.1% Tween-20) and incubated in exposure buffer (100mM Tris Base, 100mM NaCl, 50mM MgCl2, 0.1% tween, 4.5µl/ml NBT and 3.5µl/ml BCIP in 10% PVA), followed by washing in v100% EtOH (20 minutes).

## Results

### Schistosome death occurs after the PZQ-evoked shift from the mesenteries to the liver

Following PZQ exposure, *S. mansoni* shift from the mesenteric vasculature, where mature worms normally reside, to the liver [22]. We harvested worms from the mesenteries and livers of mice at various time points after PZQ (400 mg/kg) treatment in order to assess the effects of *in vivo* chemotherapy on parasites (Figure 1A). Worms were processed for measurement of pro-apoptotic caspase 3/7 activation, a readout of worm death, and imaging by transmission electron microcopy, to visualize changes to tissue ultrastructure. Worms displayed activated caspase 3/7 activity beginning 3 hours after PZQ exposure, and this signal reached a maximum at 24 hours after drug treatment. This readout of worm death occurred after the hepatic shift – which began minutes after PZQ administration and was complete within 6 hours (Figure 1B). In parallel to processing these samples, worms were harvested for RNA-Seq at timepoints shown in Figure 1B. A total of 1848 transcripts were evidenced by an average TPM >3 across the timepoints studied and displayed >1 log_2_ fold change relative to the controls t=0 timepoint (S1 File). Hierarchical clustering of these data revealed that the timepoints clustered into two groups, 0 – 9 hours and 12 – 96 hours (Figure 1C). The onset of the greatest transcriptional changes, at around 12 hours after PZQ treatment, corresponds to the period following the parasite hepatic shift.

While changes such as hepatic shift, caspase activation and gene expression took several hours, PZQ caused rapid changes to schistosome tissue ultrastructure. The parasite tegument sits a top layers of body wall muscle, which exhibit a ‘bunched’ appearance at the earliest timepoint measured after drug administration (15 minutes). However, this effect was not apparent at later timepoints. From 3 hours onward the muscle and tegument displayed a loss of integrity with extensive vacuolization. By 24 hours post drug exposure, the tegument was almost entirely missing in certain regions, exposing underlying layers of body wall muscle (Figure 1D). This time course is notable because PZQ has a relatively short half-life *in vivo*. In humans, PZQ’s half-life is approximately 2 hours (reviewed in [23]), and elimination is likely even more rapid in mice. This brief window corresponds to the changes in worm musculature observed within an hour after drug treatment, but not the window of corresponding to worm death and the most dramatic changes in gene expression. Therefore, these data focused our attention on the events that occur following the parasite hepatic shift, between 12-24 hours after drug exposure.

### Establishing sublethal conditions for *in vivo* PZQ treatment

In order to identify an active dose of PZQ for RNA-Seq studies that did not lethally and irreversibly impact schistosomes, we administered various doses of PZQ to mice harboring 7-week-old infections. As expected, treatment with a fully curative dose of PZQ (400 mg/kg) caused an irreversible hepatic shift. However, mice treated with low dose PZQ (100 mg/kg) exhibited only a transient parasite hepatic shift (Figure 2A), with worms recovering and returning to the mesenteric vasculature within two days. The sublethal effect of PZQ (100 mg/kg) was verified by measurement of pro-apoptotic caspase activity. Worms harvested from the livers of mice treated with PZQ (400 mg/kg) 14 hours after drug treatment exhibit a 71.0 ± 3.6-fold increase in caspase activation. However, worms harvested from the livers of mice treated with a low dose of PZQ (100mg/kg) at the same timepoint exhibit only a 5.0 ± 0.8-fold increase in caspase activation relative to vehicle controls (Figure 2B). Therefore, low dose PZQ (100 mg/kg) was used for subsequent transcriptomic studies given a sublethal activity on schistosomes.

**Figure 2.**
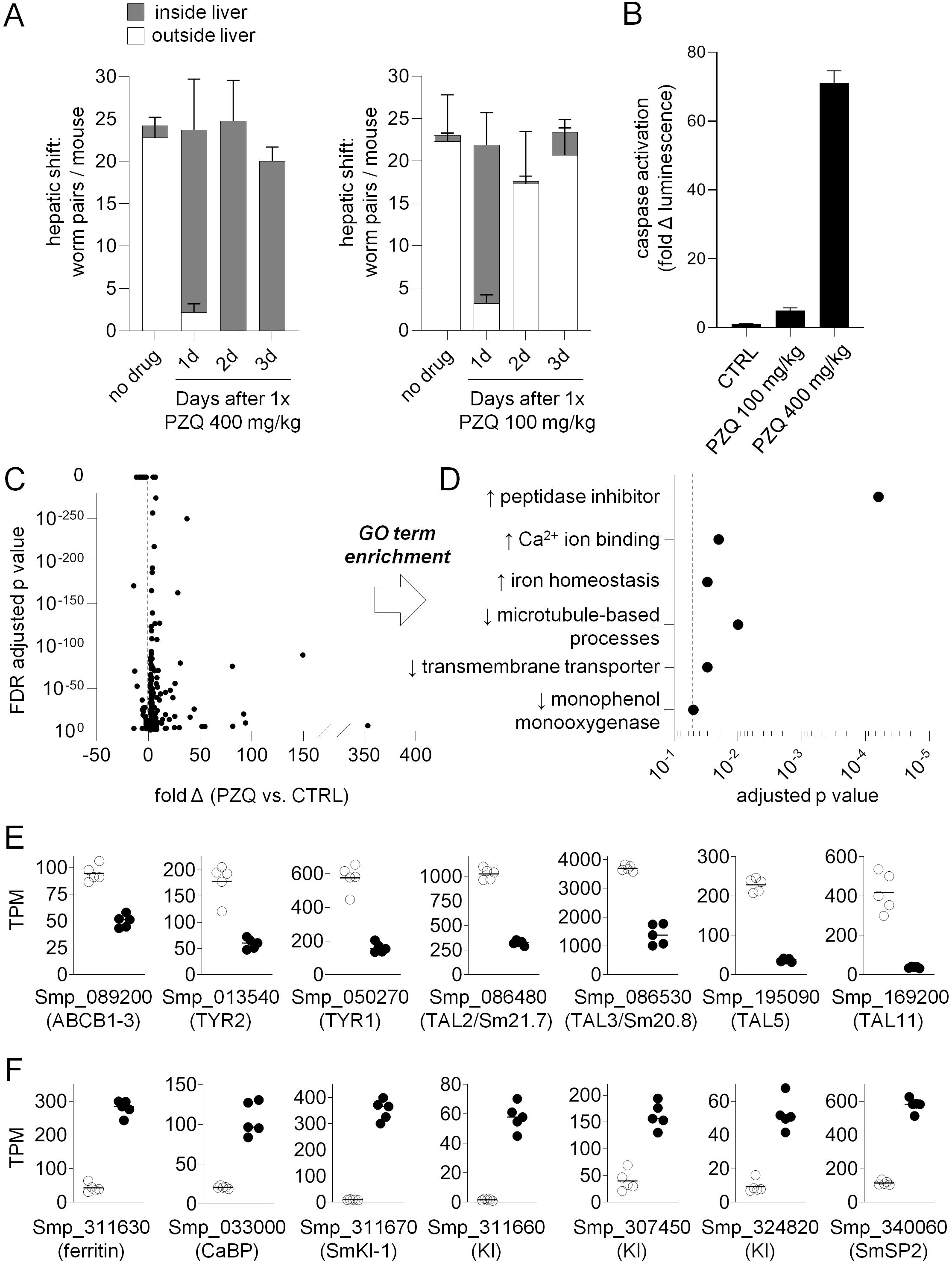
Transcriptional response of mature *S. mansoni* to a sublethal dose of praziquantel. **(A)** Mice were administered PZQ 400 mg/kg or PZQ 100 mg/kg and then euthanized at various time points to count the proportion of parasites found either within the liver (grey stacked bars) or outside the liver (white stacked bars). **(B)** Measurement of pro-apoptotic caspase-3/7 activation in homogenate of worms harvested from the livers of mice one day after treatment with PZQ 100 mg/kg or 400 mg/kg. **(C)** Volcano plot of transcripts differentially expressed between worms harvested from mice treated with PZQ (100 mg/kg) or vehicle control. **(D)** Gene-ontology (GO) term enrichment of up-regulated (↑) and down-regulated (↓) lists of transcripts (dashed line, p=0.05). **(E-F)** Examples of differentially expressed gene products containing GO-term annotations in (D). (E) PZQ down-regulated transcripts and (F) PZQ up-regulated transcripts. These data include gene products reported in prior studies (PZQ down-regulation of ABCB1-3 and tyrosinase isoforms, and PZQ up-regulation of ferritin and CaBP isoforms) as well as down-regulated tegument like allergens (TALs) and up-regulated peptidase inhibitors (Kunitz-type protease inhibitors). Symbols represent TPM (Transcripts Per Million) from parasites harvested from individual mice (n=5 independent biological replicates) treated with vehicle control (open symbols) or PZQ (solid symbols). Bar = mean TPM value for each cohort.

### Transcriptional response of mature parasites to *in vivo* PZQ exposure

Having established a sub-lethal dose of chemotherapy, we analyzed gene expression in 7-week-old parasites harvested from mice 14 hours after treatment with PZQ (100 mg/kg). Equal numbers of male and female worms were harvested from the livers of PZQ treated mice or from the mesenteries of vehicle control treated animals and processed for Illumina sequencing. Reads were mapped to the *S. mansoni* genome (v7) and differential gene expression was assessed between control and PZQ treated samples (S2 File). Up and down-regulated gene products were filtered based on ≥2-fold change, FDR adjusted p-value < 0.05, and mean expression level >2 TPM in PZQ treated samples for up-regulated transcripts and mean expression level >2 TPM in control samples for down-regulated transcripts. This revealed 201 transcripts down-regulated and 204 transcripts up-regulated with PZQ treatment (Figure 2C, S2 File).

Broadly, these data confirmed differentially expressed transcripts reported in prior microarray studies (Figure 2D-F). For example, down-regulated gene products were enriched in gene ontology (GO) terms such as transmembrane transporter activity (ex. ABC transporter ABCB1-3, Smp_089200, which decreases in *S. mansoni* following PZQ exposure [17, 24]) and monooxygenase activity (ex. Tyrosinase isoforms required for egg production [25] that are down-regulated in *S. japonicum* following PZQ treatment [18]). Up-regulated transcripts include numerous calcium ion binding proteins, although with smaller predicted molecular weights (∼8-10 kDa) than would be expected for calmodulins. These include various 8 kDa Ca^2+^ binding proteins (CaBPs) such as Smp_033000, Smp_032990, and Smp_335140 (the homolog of the PZQ up-regulated *S. japonicum* Contig10880 [18]). Ferritin isoforms (Smp_311630 & Smp_311640) were also up-regulated, as observed in [17, 24]). However, the most enriched GO term, ‘peptidase inhibitor’, was associated with a set of Kunitz-type protease inhibitors (Smp_337730, Smp_311660, Smp_311670, Smp_307450, and Smp_324820) not previously reported in other studies of PZQ response – perhaps because these gene models were recently added in the *S. mansoni* v7 genome.

These transcriptome data also reveal a caveat for utilizing GO term or pathway analysis to study schistosome datasets. These approaches rely on gene annotations mapped from better studied model organisms. However, schistosomes contain many gene products that are unique to flatworms, and either lack annotated protein domains or encode unique proteins that utilize these domains in novel ways. We found that PZQ up and down-regulated gene products were frequently unannotated and more likely to lack GO term annotations, PFAM domains, or have a BLASTp hit in well-studied model organisms (S1 Figure). Many flatworm-specific gene products have not been studied and have unknown expression patterns and function. However, several gene families found within our differentially expressed transcripts have putative roles in parasite development or host-parasite interactions [26-31]. Micro-exon gene (MEG) members are both up-regulated (Smp_336990 (MEG-2.2) and Smp_127990 (MEG-13)) and downregulated (Smp_163710 (MEG-6) and Smp_243770 (MEG-29)). Numerous egg protein CP391S-like transcripts are up-regulated (Smp_194130, Smp_102020, Smp_179970, Smp_201330), as well as several orphan lymphocyte antigen 6 (Ly6) members (transcripts Smp_105220 (SmLy6B), Smp_081900 (SmLy6C), Smp_166340 (SmLy6F), and Smp_345020 (SmLy6J)). Finally, parasite allergens were down-regulated with PZQ treatment, including venom allergen-like (VAL) transcripts (Smp_124060 (SmVAL13) and Smp_154290 (SmVAL27)) and flatworm-specific tegumental allergen-like (TAL) transcripts. These TAL gene products - Smp_086480 (SmTAL2 or Sm21.7), Smp_086530 (SmTAL3 or Sm20.8), Smp_195090 (SmTAL5), and Smp_169200 (SmTAL11) – contain dynein-light-chain domains that account for GO term enrichment related to microtubule-based processes and transport (Figure 2D).

### Transcriptional response of immature schistosomes to *in vivo* PZQ exposure

While PZQ cures infections at the sexually mature (7-week-old) parasite stage, the drug is ineffective *in vivo* against immature (4-week-old) infections (Figure 3A, [16]). However, *in vitro* PZQ treatment has similar effects on either 4-week or 7-week old parasites, causing contractile paralysis at approximately equal concentrations (Figure 3B, [32]). Instead, the major difference between these two developmental stages appears to be the effect of PZQ treatment on neoblast-like, mitotically active cells. Immature 4-week-old worms exposed to PZQ for 12 hours, followed by a pulse of EdU, show retained mitotic activity - even after PZQ doses as high as 10 μM. However, similarly treated mature 7- week old worms display at loss of mitotic activity following treatment with concentrations of PZQ ranging from 0.1 – 0.5 μM (Figure 3C).

**Figure 3.**
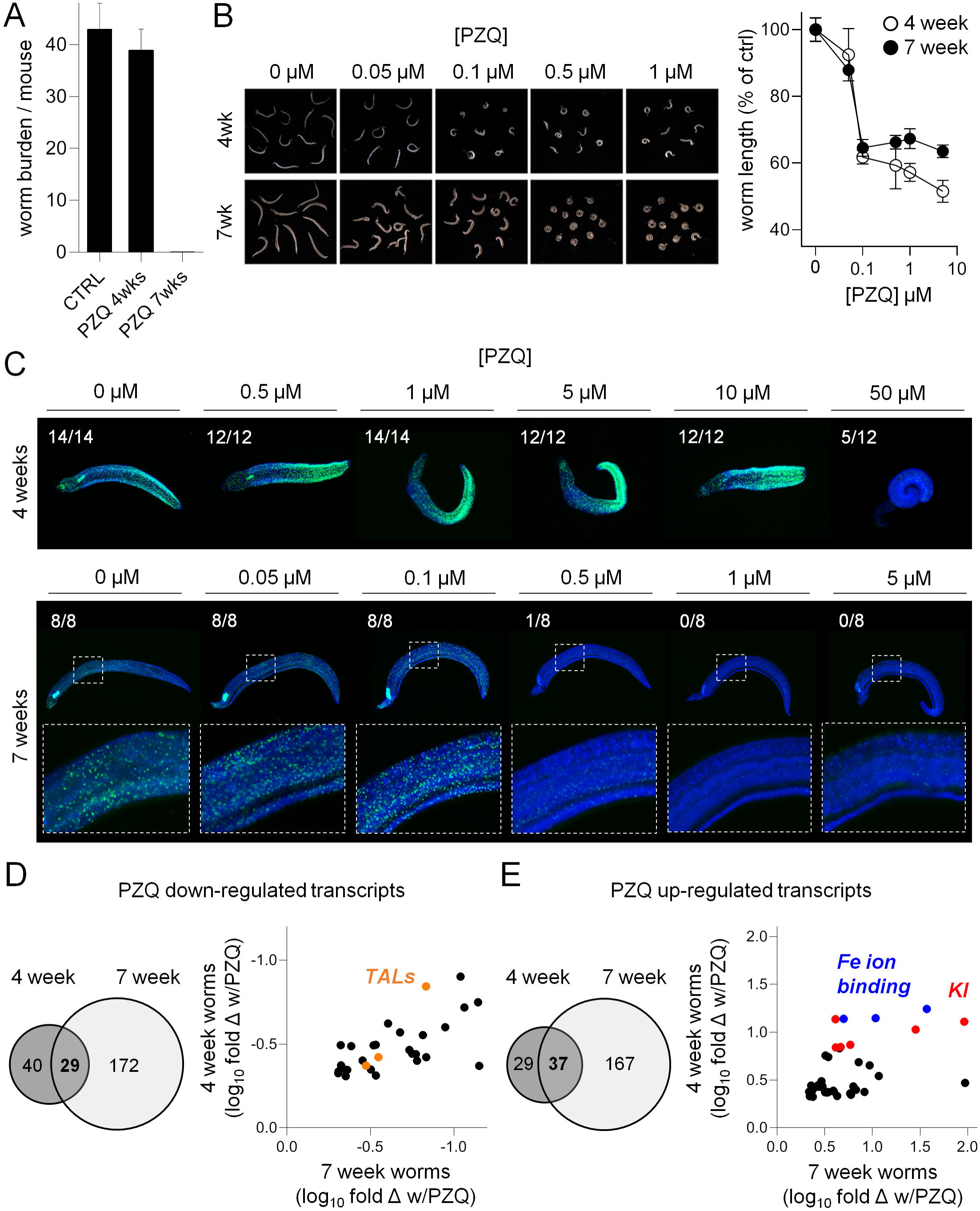
Praziquantel-evoked transcriptional changes in immature schistosomes. **(A)** Immature, 4-week-old parasites are unresponsive to PZQ treatment *in vivo*, while 7-week-old infections are cleared. However, **(B)** PZQ has comparable effects on the contraction of both immature and mature worms treated *in vitro*. Left = images of worms treated with varying concentrations of PZQ. Right = quantification of worm body length as a measure of contractile paralysis. **(C)** Effects of *in vitro* PZQ treatment (14 hours) on the mitotic activity of 4-week and 7-week-old worms (worms harvested from mice at 25 and 49 days post-infection, respectively, plus 2 days in culture). Green = EdU incorporation. Blue = DAPI counterstain. Scoring reflects number of EdU positive worms per treatment condition. **(D-E)** Venn diagram of down-regulated and up-regulated transcripts following PZQ treatment of 4-week-old and 7-week-old worms. Scatter plots reflect log_10_ fold change of transcripts found in the intersection of both datasets relative to vehicle control. Orange = tegument like allergens (TALs). Red = Kunitz-type protease inhibitors (KI). Blue = Iron ion binding gene products.

Given these data, we were interested to see to what extent the transcriptional response of 4- week-old worms to *in vivo* PZQ exposure resembled that of 7-week-old worms. Mice were dosed with PZQ (100 mg/kg) or vehicle control 4 weeks post-infection, euthanized 14 hours later, and parasites were harvested from the livers for comparative RNA-Seq just as for 7-week old samples. There were less differentially regulated transcripts in 4-week-old worms relative to the 7-week dataset (69 PZQ down-regulated transcripts and 66 PZQ up-regulated transcripts, S3 File). Of the transcripts differentially expressed in immature worms with PZQ treatment, roughly half were found in the 7-week dataset. GO term enrichment in the 4-week-old worm dataset was broadly similar to the 7-week dataset (S1 Table). PZQ down-regulated gene products in each dataset included various TAL gene products (SmTAL2, SmTAL3 and SmTAL5, Figure 3D), and PZQ up-regulated gene products in both 4 and 7-week old worms included Kunitz-type protease inhibitors and ferritin isoforms (Figure 3E).

### Tissue localization of transcripts differentially expressed following PZQ exposure

We performed *in situ* hybridization to localize the expression patterns of PZQ up and down-regulated gene products in adult, 7-week-old worms. Many down-regulated transcripts localized to the germ line (Figure 4A) – with expression patterns staining the female vitellaria (Smp_076320 (myb/sant-like) and ovaries (Smp_246770 (cadherin)), as well as the male testes (Smp_195090 (SmTAL5)). This is consistent with PZQ treatment causing loss of mitotic activity in germ line tissues.

**Figure 4.**
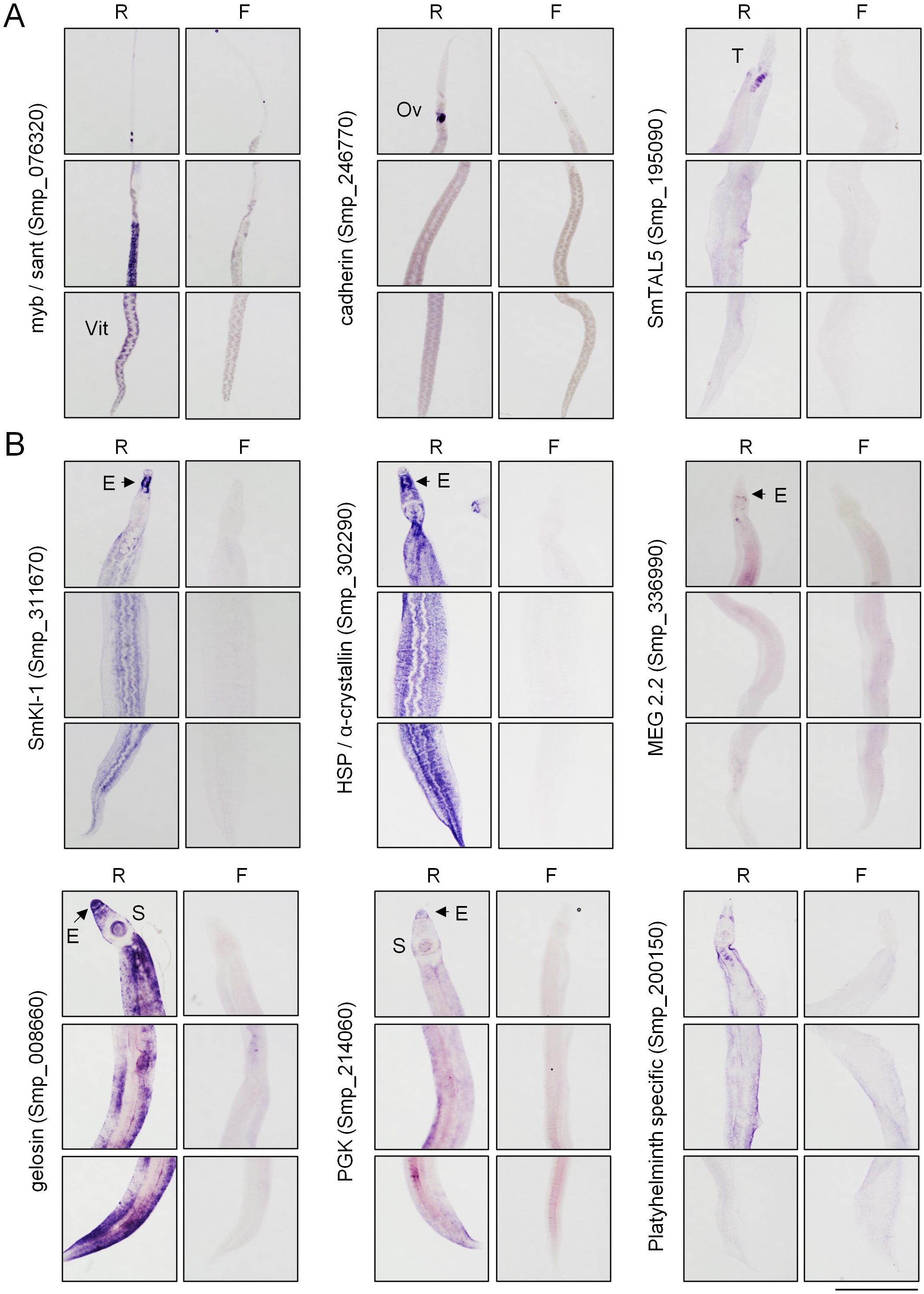
Expression patterns of transcripts differentially regulated with praziquantel treatment. *In situ* hybridization of transcripts **(A)** down-regulated and **(B)** up-regulated following *in vivo* PZQ treatment relative to vehicle controls. F = sense negative control probes. R = antisense probes. Images show, from top to bottom, anterior to posterior panels of worms. Ov = ovaries, T = testis, Vit = vitellaria, E = esophagus, S = oral sucker. Scale = 1mm.

Many PZQ up-regulated transcripts, such as Kunitz-type protease inhibitors, heat-shock protein, MEG 2.2, alpha-crystallin and phosphoglycerate kinase displayed expression patterns with varied localization within the male body. However, these commonly displayed strong expression at the anterior of the worm, with staining glands located around the esophagus (Figure 4B). The schistosome esophagus has been shown to be a secretory organ [33], and various MEG and VAL gene products have been localized to this structure [27].

### Time course of PZQ-evoked changes in gene expression

Given that our RNA-Seq was generated on worms harvested at a relatively late timepoint after drug exposure, we wanted to establish whether these data reflect acute, drug-evoked transcriptional changes or a response to later events such as the parasite hepatic shift. Therefore, we administered a single dose of PZQ (400 mg/kg) to mice harboring 7-week old infections and harvested parasites at various timepoints (from 15 minutes to 4 days) for analysis of gene expression. These data revealed that the observed changes in gene expression, such as up-regulation of Kunitz-type protease inhibitors and down-regulation of TALs, occurred relatively late after drug administration, rather than within the short window during which PZQ reaches a C_max_ *in vivo* (Figure 5A). These timepoints correspond to the parasite hepatic shift (∼12 hours onward), and may reflect the parasite response to the change in location within the host, as worms shift from the mesenteric vasculature to the Th2 environment of the granulomatous liver rich in macrophages and granulocytes (Figure 5B).

**Figure 5.**
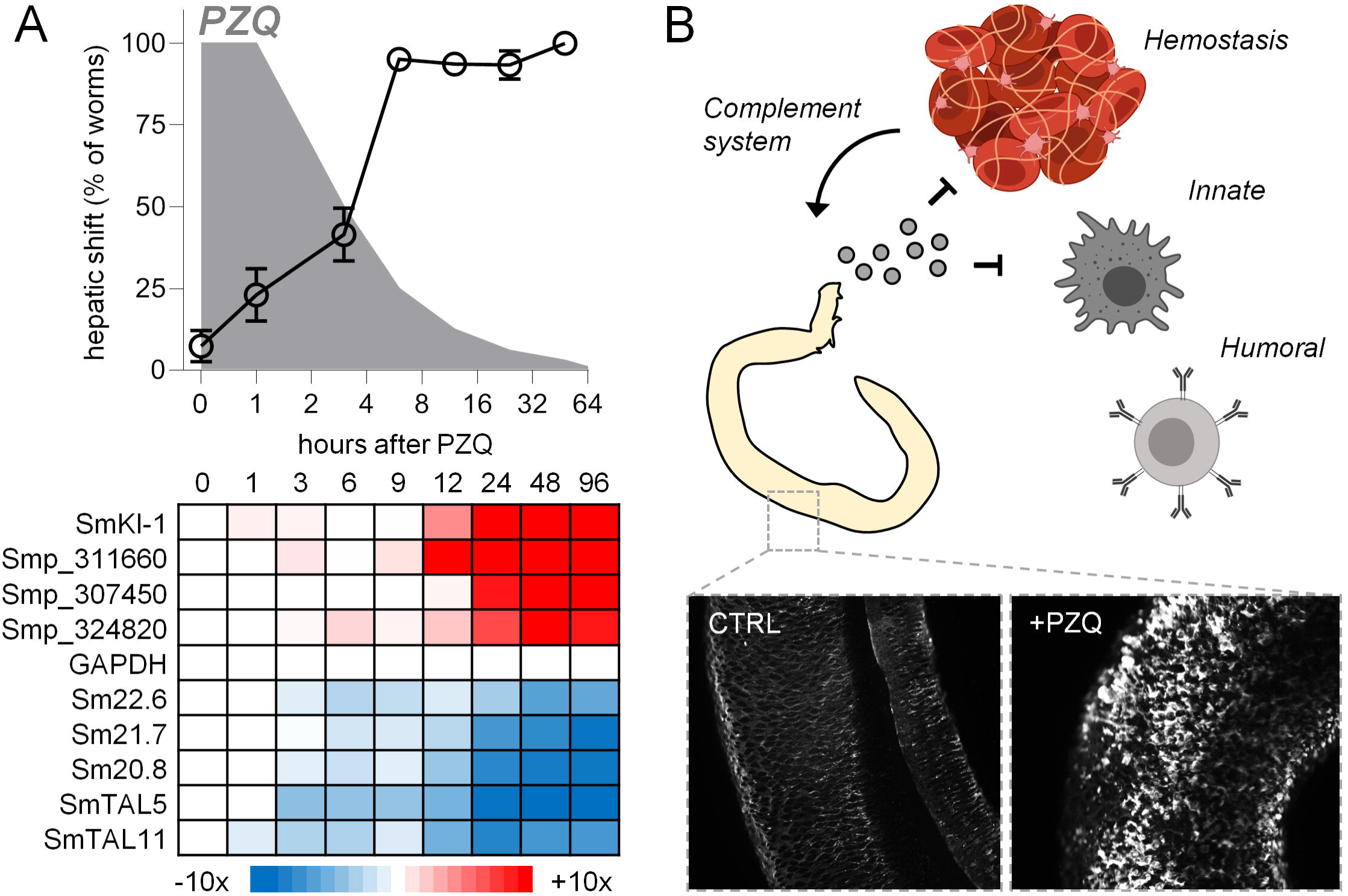
PZQ-evoked changes in immunomodulatory gene products corresponds to the onset of the parasite hepatic shift. **(A)** *Top* - Kinetics of parasite hepatic shift following PZQ (open symbols, data from Figure 1B) and predicted PZQ elimination (grey). *Bottom* - Expression (fold change) of various immunomodulatory gene products such as Kunitz-type protease inhibitors (increasing, red) and tegument-like allergens (decreasing, blue) in worms harvested at various points in the PZQ time-course shown in Figure 1. (B) Model for schistosome release of immunomodulatory signals in response to chemotherapy. Worms normally reside within the host circulatory system, evading detection by the innate and humoral immune system. The parasite tegument is damaged following PZQ exposure, and worms are exposed to the milieu of immune cells within the liver. Secreted signals may dampen the immune response, as well as impair coagulation and resulting activation of the complement pathway. Images created with BioRender.com.

## Discussion

While PZQ has been the frontline anthelmintic used to control schistosomiasis for over 40 years, the drug’s molecular mechanism of action is poorly understood. From *in vitro* studies it is clear that PZQ has pronounced effects on parasite musculature and tegument [5, 32]. However, we were interested in several apparent inconsistencies between *in vitro* and *in vivo* observations of PZQ activity. First, it is not clear that PZQ is directly schistocidal in vivo. That is, while *in vitro* experiments often measure worm death after periods of drug incubation, PZQ has a short half-life *in vivo* (∼2 hours in humans, reviewed in [23]). Worms harvested from mice after treatment with PZQ indeed display rapid changes in muscle structure (within minutes of drug administration, Figure 1D). However, this effect was transient, and outcomes such as parasite tegument damage, broad transcriptional changes and death did not occur until hours later - reaching a maximum at one day after drug treatment. Second, PZQ does not cure immature 4-week-old infections [34, 35]. This is a clinically important feature of PZQ that may underpin treatment failure in areas of high transmission [36]. However, it is not entirely accurate to say that immature worms are unresponsive to PZQ, since the drug causes contractile paralysis of both 4-week and 7-week-old parasites *in vitro* with approximately equal potency ([32], Figure 3B). Therefore, in order to better understand the effect of *in vivo* PZQ exposure on *S. mansoni*, we performed comparative RNA-Seq on mature and immature worms harvested from PZQ-treated mice. These data provide an overview of not just direct PZQ-evoked changes in gene expression (as may be the case with *in vitro* PZQ treatment [17]), but also the worm response to enviornmental change (i.e. shift from the mesenteric vasucalture to the liver) and attack by components of the host immune system.

### Comparative responses of immature and mature schistosomes to PZQ

The transcriptional response to *in vivo* PZQ exposure is similar between 4-week-old and 7-week-old parasites (Figure 3, S1 Table) – although mature parasites show greater changes in gene expression, perhaps reflecting a greater sensitivity to chemotherapy. It has also been speculated that lack of in vivo PZQ efficacy against 4-week-old parasites may be due to PZQ pharmacokinetics, since mature worms within the mesenteric vasculature are exposed to higher drug concentrations prior to first pass metabolism [23, 37]. Another possibility is the difference in the host immune environment at the 4- week stage of infection relative to mature infections [38, 39]. Indeed, both these factors may contribute to a lack of PZQ efficacy against juvenile worms. However, the neoblast-like cells of immature and mature worms are affected differently following *in vitro* PZQ exposure (Figure 3C), indicating that there are inherent differences between these stages. Since immature worms harbor more abundant stem cells, such as transitory somatic ε-cells [40], this may account for treatment failure during these developmental stages.

### PZQ-evoked changes in immunomodulatory gene products

Many of the gene products differentially regulated by PZQ modulate the host immune system. What might be the biological effect of PZQ regulation of immunomodulatory gene products? Various blood-dwelling parasitic helminths secrete immunomodulatory vesicles into the host circulation [3, 41, 42], and under normal infection conditions schistosomes modulate components of the host circulatory system. For example, blood from mice harboring patent schistosome infections displays altered clotting properties relative to uninfected mice or mice with immature infections [43]. Parasites may up-regulate anti-clotting signals as an immune-evasion mechanism, since fibrin clots serve as a scaffold for adhesion of granulocytes and monocytes, and activated platelets regulate recruitment and actions of innate immune cells (reviewed in [44]). Schistosomes are susceptible to attack by the host complement system (reviewed in [45]), which is activated by enzymes in the coagulation cascade such as FXa. Therefore, up-regulation of anti-coagulant gene products may enable schistosomes to survive transient PZQ exposure *in vivo* and ultimately resume patent infections within the mesenteric vasculature.

Many immunomodulatory gene products are expressed in the schistosome esophagus [33]. These gene products are likely important internally, protecting the parasite from ingested immune components and enzymes found in leukocytes and erythrocytes [27, 46-49], and they may also be secreted outside of the worm to modulate various immune cells within the host circulation. PZQ up-regulation of esophageal immunomodulatory gene products may be a parasite response to evade recognition by the host immune system, triggered either by drug-evoked tegument damage or the hostile immune environment of the liver.

For example, the most enriched group of up-regulated transcripts were Kunitz-type protease inhibitors. One of these, SmKI-1, has been characterized and shown to inhibit neutrophil function [19] and impair blood coagulation via inhibition of enzymes such as FXa [50]. *In situ* hybridization localized PZQ up-regulated Kunitz-type protease inhibitors to the schistosome esophagus (Figure 4B). Blood-feeding animals harbor various Kunitz-type protease inhibitors with anti-coagulant activity [51] and these proteins are enriched in the salivary proteomes of these organisms [52, 53]. Kunitz-type protease inhibitors have also been found in the secretomes of other parasitic flatworms [54, 55], and shown to act as ion channel blockers [56] and inhibitors of dendritic cell activation [57]. Additionally, laboratory strains of *S. mansoni* selected for PZQ resistance show altered expression of a various Kunitz-type protease inhibitors, indicating these may be involved in drug resistance [58].

While secreted proteins may promote immune evasion in the short term, numerous gene products in our PZQ up-regulated dataset have also been proposed as schistosomiasis vaccine targets. This includes SmKI-1 [59, 60], but also cathepsins [61], MEGs [62] and tetraspanins [63]. Therefore, secreted signals may assist in evasion of the innate immune system while also promoting development of host antibodies against parasite antigens. This acquired immunity may not be deleterious to existing schistosomes, which are able to survive within the circulatory system alongside host antibodies and immune cells [64], but recognition of parasite antigens may confer immunity to new infections following chemotherapy [65-67].

These data are the first comparative RNA-Seq dataset on *S. mansoni* exposed to PZQ in vivo. Our findings confirm changes in gene expression reported in prior *in vitro* studies and microarray experiments, as well as revealing changes in the expression of immunomodulatory gene products that localize to the parasite esophageal glands. Given that several of these gene products, such as the Kunitz-type protease inhibitors, have anti-coagulant effects in both schistosomes and other blood feeding parasites and vectors, these changes may reflect a mechanism employed by schistosomes to actively subvert the hemostatic system and evade the host immune system in response to chemotherapy. These mechanisms inform our basic understanding of parasite interaction with their hosts and provide insight into potential mechanisms for PZQ treatment failure or routes to anthelmintic drug resistance.

## Acknowledgements

The following reagent was provided by the NIAID Schistosomiasis Resource Center for distribution through BEI Resources, NIH-NIAID Contract HHSN272201700014I. NIH: *Schistosoma mansoni*, Strain NMRI, Exposed Swiss Webster Mice, NR-21963.

## Supporting Information

**S1 Figure. Parasite-specific gene products are differentially expressed in praziquantel treated worms.** X-axis = Gene products evidenced by read mapping >0 ranked from most up-regulated to most down-regulated following PZQ treatment. Y-axis = number of transcripts that lack a GO term annotation (top), PFAM protein domain (middle) or BLASTp hit verses the landmark database (bottom) for every 100 gene products.

**S1 Table. GO-term enrichment in differentially expressed transcripts following PZQ treatment.** GO-term enrichment from list up up-regulated and down-regulated transcripts (at least two-fold change, FDR adjusted p value < 0.05) for both immature (4-week) and mature (7-week) infections. n.s. = not significant.

**S1 File. Time course RNA-Seq data following *in vivo* PZQ treatment. (Sheet 1)** Read counts or **(Sheet 2)** Transcripts Per Million (TPM) for transcripts in worms harvested from mice treated with PZQ (400mg/kg). **(Sheet 3)** Z-scores of 1848 transcripts with an average TPM >3 and >1 log_2_ fold change relative to the t=0 timepoint that were used to generate the heat map in Figure 1C.

**S2 File. RNA-Seq data for 7-week worms following *in vivo* PZQ treatment. (Sheet 1)** Read counts or **(Sheet 2)** Transcripts Per Million (TPM) for transcripts in worms harvested from mice (n=5) treated with either vehicle control or PZQ (100mg/kg) 7-weeks post-infection. **(Sheet 3)** List of filtered PZQ up and down-regulated transcripts.

**S3 File. RNA-Seq data for 4-week worms following *in vivo* PZQ treatment. (Sheet 1)** Read counts or **(Sheet 2)** Transcripts Per Million (TPM) for transcripts in worms harvested from mice (n=5) treated with either vehicle control or PZQ (100mg/kg) 4-weeks post-infection. **(Sheet 3)** List of filtered PZQ up and down-regulated transcripts.

**S4 File. Primers used in generation of *in situ* hybridization probes.** Primer sequences for forward and reverse *in situ* hybridization probes used in Figure 4.

