## Supplementary figures and images for "Schistosomes alter expression of immunomodulatory gene products following *in vivo* praziquantel exposure"

### Supplemental Figure 1

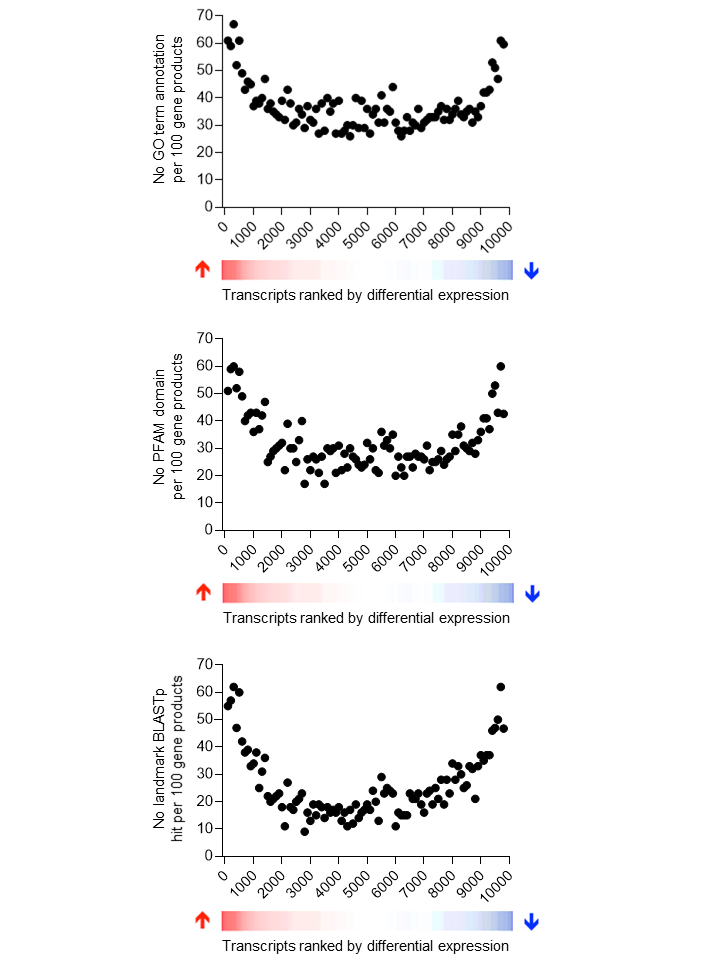
