## Supplemental Table 1 for "Schistosomes alter expression of immunomodulatory gene products following *in vivo* praziquantel exposure"

|  | **Source** | **Term name** | **Term ID** | **Immature worms**  **(adj. p value)** | **Mature worms**  **(adj. p value)** |
| --- | --- | --- | --- | --- | --- |
| PZQ  up-regulated | GO:MF | peptidase inhibitor activity | GO:0030414 | 2.50E-06 | 6.40E-05 |
|  | GO:MF | phospholipid binding | GO:0005543 | 1.80E-03 | n.s. |
|  | GO:MF | calcium ion binding | GO:0005509 | 2.00E-02 | n.s. |
|  | GO:MF | iron ion binding | GO:0005506 | 3.80E-02 | 3.40E-02 |
| PZQ  down-regulated | GO:BP | microtubule-based process | GO:0007017 | 8.00E-04 | 9.60E-03 |
|  | GO:MF | extracellular matrix structural constituent | GO:0005201 | 8.71E-07 | n.s. |
|  | GO:MF | active transmembrane transporter activity | GO:0022804 | n.s. | 3.10E-02 |
|  | GO:MF | monophenol monooxygenase activity | GO:0004503 | n.s. | 5.00E-02 |
